# A Bayesian Framework for Estimating Cell Type Composition from DNA Methylation Without the Need for Methylation Reference

**DOI:** 10.1101/112417

**Authors:** Elior Rahmani, Regev Schweiger, Liat Shenhav, Theodora Wingert, Ira Hofer, Eilon Gabel, Eleazar Eskin, Eran Halperin

## Abstract

We introduce a Bayesian semi-supervised method for estimating cell counts from DNA methylation by leveraging an easily obtainable prior knowledge on the cell type composition distribution of the studied tissue. We show mathematically and empirically that alternative methods which attempt to infer explicit cell counts without methylation reference can only capture linear combinations of cell counts rather than provide one component per cell type. Our approach, which allows the construction of a set of components such that each component corresponds to a single cell type, therefore provides a new opportunity to investigate cell compositions in genomic studies of tissues for which it was not possible before.

## 1 Background

DNA methylation status has become a prominent epigenetic marker in genomic studies, and genome-wide DNA methylation data have become ubiquitous in the last few years. Numerous recent studies provide evidence for the role of DNA methylation in cellular processes and in disease (e.g., in multiple sclerosis [1], schizophrenia [2], and type 2 diabetes [3]). Thus, DNA methylation status holds great potential for better understanding the role of epigenetics, potentially leading to better clinical tools for diagnosing and treating patients.

In a typical DNA methylation study, we obtain a large matrix in which each entry corresponds to a methylation level (a number between 0 and 1) at a specific genomic position for a specific individual. This level is the fraction of the probed DNA molecules that were found to have an additional methyl group at the specific position for the specific individual. Essentially, these methylation levels represent, for each individual and for each site, the probability of a given DNA molecule to be methylated. While simple in principle, methylation data are typically complicated owing to various biological and non-biological sources of variation. Particularly, methylation patterns are known to differ between different tissues and between different cell types. As a result, when methylation levels are collected from a complex tissue (e.g., blood), the observed methylation levels collected from an individual reflect a mixture of its methylation signals coming from different cell types, weighted according to mixing proportions that depend on the individual’s cell type composition. Thus, it is challenging to interpret methylation signals coming from heterogeneous sources.

One notable challenge in working with heterogeneous methylation levels has been highlighted in the context of Epigenome-Wide Association Studies (EWAS), where data are typically collected from heterogeneous samples. In such studies, we typically search for rows of the methylation matrix (each corresponding to one genomic position) that are significantly correlated with a phenotype of interest across the samples in the data. In this case, unless accounted for, correlation of the phenotype of interest with the cell type composition of the samples may lead to numerous spurious associations and potentially mask true signal [4]. In addition to its importance for a correct statistical analysis, knowledge of the cell type composition may provide novel biological insights by studying cell compositions across populations.

In principle, one can use high-resolution cell counting for obtaining knowledge about the cell composition of the samples in a study. However, unfortunately, such cell counting for a large cohort may be costly and often logistically impractical (e.g., in some tissues, such as blood, reliable cell counting can be obtained from fresh samples only). Due to the pressing need to overcome this limitation, development of computational methods for estimating cell type composition from methylation data has become a key interest in epigenetic studies. Several such methods have been suggested in the past few years [5, 6, 7, 8, 9, 10], some of which aim at explicitly estimating cell type composition, while others aim at a more specific goal of correcting methylation data for the potential cell type composition confounder in association studies. These methods take either a supervised approach, in which reference data of methylation patterns from sorted cells (methylomes) are obtained and used for predicting cell compositions [5], or an unsupervised approach (reference-free) [6, 7, 8, 9, 10].

The main advantage of the reference-based method is that it provides direct (absolute) estimates of the cell counts, whereas, as we demonstrate here, current reference-free methods are only capable of inferring components that capture linear combinations of the cell counts. Yet, the reference-based method can only be applied when relevant reference data exist. Currently, reference data only exist for blood [11], breast [12] and brain [13], for a small number of individuals (e.g., six samples in the blood reference [11]). Moreover, the individuals in most available data sets do not match the reference individuals in their methylation-altering factors, such as age [14], gender [15, 16], and genetics [17]. This problem was recently highlighted in a study in which the authors showed that available blood reference collected from adults failed to estimate cell proportions of newborns [18]. Furthermore, in a recent work, we showed evidence from multiple data sets that a reference-free approach can provide substantially better correction for cell composition when compared with the reference-based method [19]. It is therefore often the case that unsupervised methods are either the only option or are a better option for the analysis of EWAS.

As opposed to the reference-based approach, although can be applied for any tissue in principle, the reference-free methods do not provide direct estimates of the cell type proportions. Previously proposed reference-free methods allow us to infer a set of components, or general axes, which were shown to compose linear combinations of the cell type composition [8, 9]. Another more recent reference-free method was designed to infer cell type proportions, however, as we show here, it only provides components that compose linear combinations of the cell type composition rather than direct estimates [10]. Unlike cell proportions, while linearly correlated components are useful in linear analyses such as linear regression, they cannot be used in any nonlinear downstream analysis or for studying individual cell types (e.g., studying alterations in cell composition across conditions or populations). Cell proportions may provide novel biological insights and contribute to our understanding of disease biology, and we therefore need targeted methods that are practical and low in cost for estimating cell counts.

In attempt to address the limitations of previous reference-free methods and to provide cell count estimates rather than linear combinations of the cell counts, we propose an alternative Bayesian strategy that utilizes prior knowledge about the cell type composition of the studied tissue. We present a semi-supervised method, BayesCCE (Bayesian Cell Count Estimation), which encodes experimentally obtained cell count information as a prior on the distribution of the cell type composition in the data. As we demonstrate here, the required prior is substantially easier to obtain compared with standard reference data from sorted cells. We can estimate this prior from general cell counts collected in previous studies, without the need for corresponding methylation data or any other genomic data.

We evaluate our method using four large methylation data sets and simulated data, and show that our method produces a set of components that can be used as cell count estimates. We observe that each component of BayesCCE can be regarded as corresponding to scaled values of a single cell type (i.e. high absolute correlation with one cell type, but not necessarily good estimates in absolute terms). We find that BayesCCE provides a substantial improvement in correlation with the cell counts over existing reference free methods (in some cases a 50% improvement). We also consider the case where both methylation and cell count information are available for a small subset of the individuals in the sample, or for a group of individuals from external data. Notably, existing reference-based and reference-free methods for cell type estimation completely ignore this potential information. In contrast, our method is flexible and allows to incorporate such information. Specifically, we show that our proposed Bayesian model can leverage such additional information for imputing missing cell counts in absolute terms. Testing this scenario on both real and simulated data, we find that measuring cell counts for a small group of samples (a couple of dozens) can lead to a further significant increase in the correlation of BayesCCE’s components with the cell counts.

## 2 Results

### Benchmarking existing reference-free methods for capturing cell type composition

We first demonstrate that existing reference-free methods can infer components that are correlated with the tissue composition of DNA methylation data collected from heterogeneous sources. For this experiment, as well as for the rest of the experiments in this paper, we used four large publicly available whole-blood methylation data sets: a data set by Hannum et al. [20] (*n* = 650), a data set by Liu et al. [21] (*n* = 658), and two data sets by Hannon et al. [22] (*n* = 638 and *n* = 665; denote Hannon et al. I and Hannon et al. II, respectively). In addition, we simulated data based on a reference data set of methylation levels from sorted leukocytes cells [11] (see Methods). While cell counts were known for each sample in the simulated data, cell counts were not available for the real data sets. We therefore estimated the cell type proportions of six major blood cell types (granulocytes, monocytes and four subtypes of lymphocytes: CD4+, CD8+, B cells and natural killer cells) based on a reference-based method [5], which was shown to reasonably estimate leukocyte cell proportions from whole blood methylation data collected from adult individuals [23, 18, 24]. Due to the absence of large publicly available data sets with measured cell counts, these estimates were considered as the ground truth for evaluating the performance of the different methods.

For benchmarking performance of existing methods, we considered three reference-free methods, all of which were shown to generate components that capture cell type composition information from methylation: ReFACTor [8], Non-Negative Matrix Factorization (NNMF) [9] and MeDeCom [10]. We evaluated six components of each of these methods - six being the number of estimated cell types composing the ground truth. We found all methods to capture a large portion of the cell composition information in all data sets; particularly, we observed that ReFACTor performed considerably better than NNMF and MeDeCom in all occasions (Supplementary Figure ??).

In spite of the fact that all three methods can capture a large portion of the cell composition variation, each component provided by these methods is a linear combination of the cell types in the data rather than an estimate of the proportions of a single cell type. As a result, as we show next, in general, these methods perform poorly when their components are considered as estimates of cell type proportions. ReFACTor was not designed for estimating cell proportions but rather for providing orthogonal principal components of the data that together capture variation in cell compositions. In contrast, NNMF and MeDeCom, which extends the underlying model in NNMF, were designed to provide estimates of cell type proportions. In addition to empirical support from the data, as we report next, we also provide a mathematical proof for the non-identifiability nature of the NMMF model, which drives solutions towards undesired linear combinations of cell type proportions rather than direct estimates of cell type proportions (see Methods).

### BayesCCE: A Bayesian semi-supervised approach for capturing cell type composition

Every method that has been developed so far for capturing cell composition signal from methylation can be classified as either reference-based, wherein a reference of methylation patterns of sorted cells is used, or reference-free, wherein cell composition information is inferred in an unsupervised manner. Our proposed method, BayesCCE, combines elements from the underlying models of previous reference-free methods with further assumptions, which together direct the solution towards inference of one component per cell type. BayesCCE does not use standard reference data of sorted methylation levels, but rather it leverages a relatively weak prior information about the distribution of cell type composition in the studied tissue. BayesCCE is fully described in the Methods section.

In order to evaluate BayesCCE we obtained prior information about the distribution of leukocyte cell type proportions in blood using high resolution blood cell counts that were previously measured in 595 adult individuals (see Methods). In concordance with the estimated cell type proportions used as the ground truth, we first considered the assumption of six constituting cell types in blood tissue (*k* = 6). We applied BayesCCE on each of the four data sets, and evaluated the resulted components. We observed that each time BayesCCE produced a set of six components such that each component was correlated with one of the cell types, as desired (Figure 1 and Supplementary Tables ?? and ??). Specifically, we found the mean absolute correlation values across all six cell types to be 0.58, 0.63, 0.45 and 0.45 in the Hannum et al., Liu et al., Hannon et al. I and Hannon et al. II data sets, respectively. We note, however, that the assignment of components into corresponding cell types could not be automatically determined by BayesCCE. In addition, in general, the BayesCCE components were not in the right scale of their corresponding cell types (i.e. each component represented the proportions of one cell type up to a multiplicative constant and addition of a constant). These symptoms are expected due to the nature of the prior information used by BayesCCE. For more details about the assignment of components into cell types and evaluation measurements see Methods.

**Figure 1:**
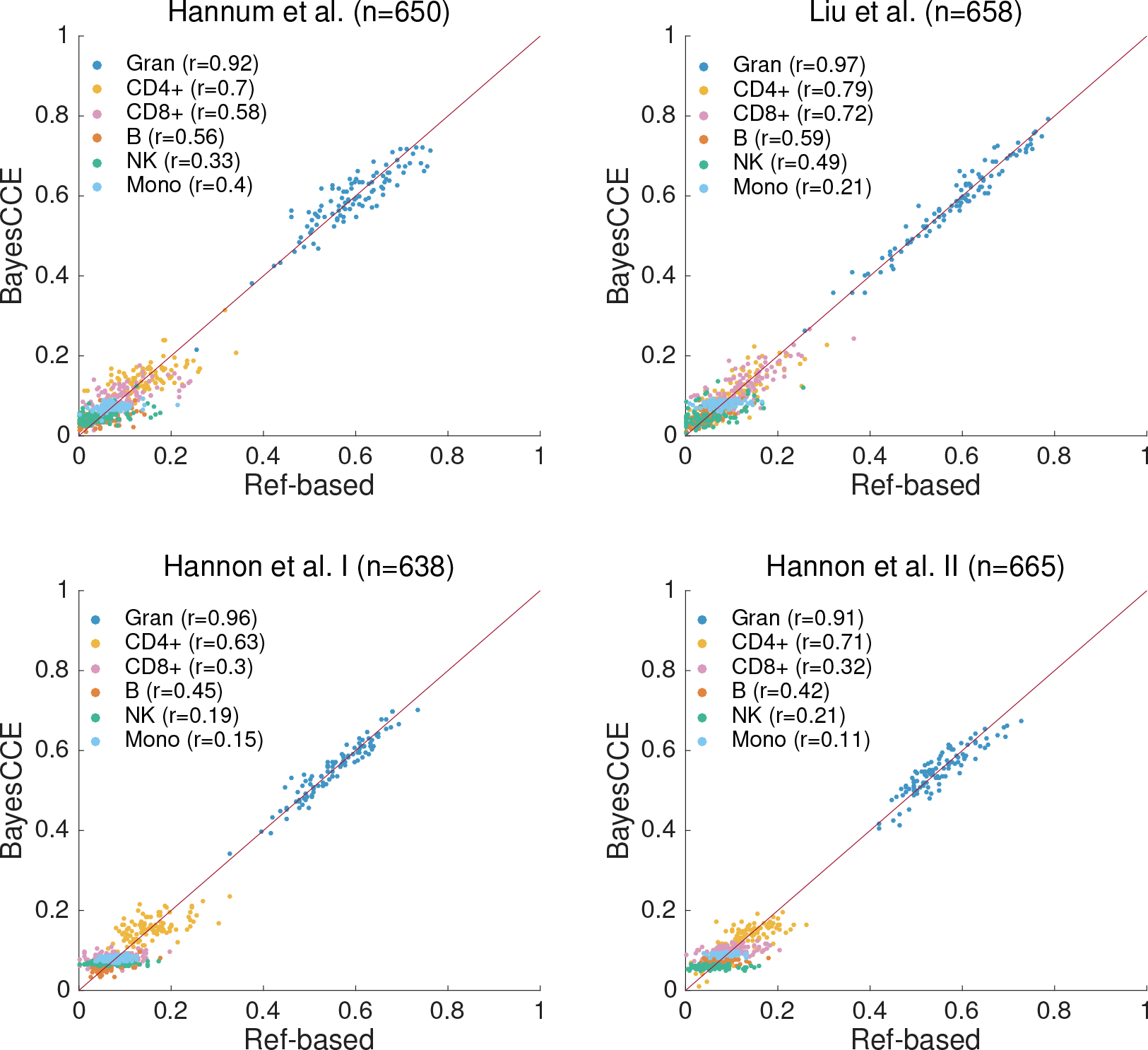
BayesCCE captures cell type proportions in four data sets under the assumption of six constituting cell types in blood (*k* = 6): granulocytes, monocytes and four subtypes of lymphocytes (CD4+, CD8+, B cells and NK cells). The BayesCCE estimated components were linearly transformed to match their corresponding cell types in scale (see Methods). For convenience of visualization, we only plot the results of 100 randomly selected samples for each data set.

We next considered a simplifying assumption of only three constituting cell types in blood tissue (*k* = 3): granulocytes, lymphocytes and monocytes. We applied BayesCCE on each of the four data sets, and observed high correlations between the estimated components of granulocytes and the granulocytes levels (*r* ≥ 0.91 in all data sets) and between the estimated components of lymphocytes and the lymphocytes levels (*r* ≥ 0.87 in all data sets), yet much lower correlations for monocytes (*r* ≤ 0.27 in all data sets; Supplementary Figure ?? and Supplementary Tables ?? and ??). We note that poor performance in capturing some cell type may be partially derived by inaccuracies introduced by the reference-based estimates, which are used as the ground truth in our experiments. Notably, three recent studies, which consisted of samples for which both methylation levels and cell count measurements were available, demonstrated that while the reference-based estimates of the overall lymphocyte and granulocyte levels were found to be highly correlated with the true levels, the accuracy of estimated monocytes was found to be substantially lower [18, 8, 25]. This may explain the low correlations we report for monocytes in our experiments. Low correlations with some of the cell types may be driven by various reasons, such as utilizing inappropriate reference or failing to perform a good feature selection. We later provide a more detailed discussion about these issues.

For assessing the performance of BayesCCE in light of previous reference-free methods, we sub-sampled the data and generated ten data sets of 300 randomly selected samples from each one of the four data sets. In addition, we simulated ten data sets of similar size (*n* = 300; see Methods). Figure 2 demonstrates a significant and substantial improvement in performance for BayesCCE upon existing methods under the assumption of six constituting cell types (*k* = 6). Repeating the same set of experiments while assuming three constituting cell types (*k* = 3) revealed similar results (Supplementary Figure ??).

**Figure 2:**
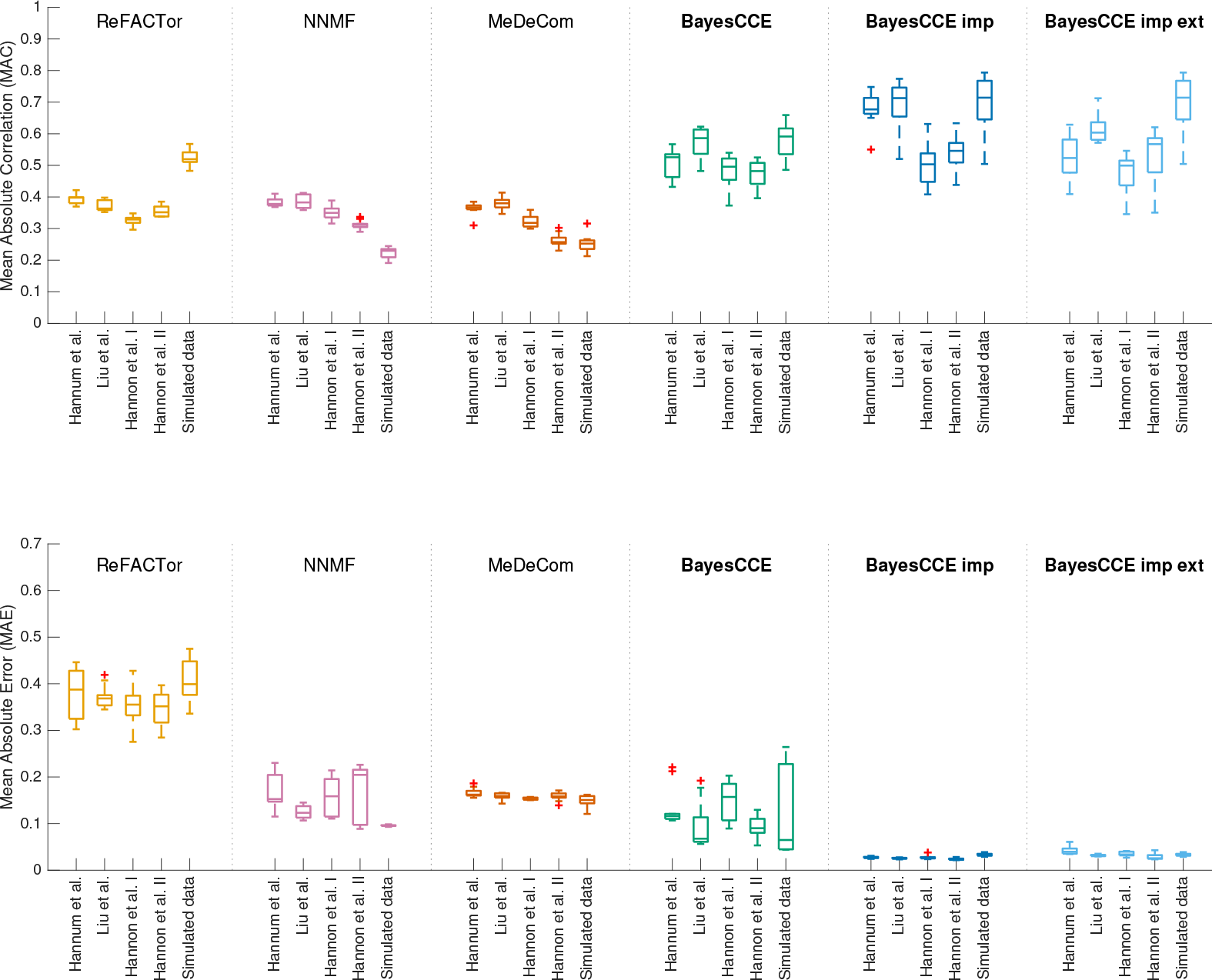
The performance of existing reference-free methods and BayesCCE under the assumption of six constituting cell types in blood (*k* = 6): granulocytes, monocytes and four subtypes of lymphocytes (CD4+,CD8+, B cells and NK cells). For each method, box plots show for each data set the performance across ten sub-sampled data sets (*n* = 300), with the median indicated by a horizontal line. For each of the methods, ReFACTor, NNMF, MeDeCom and BayesCCE, we considered a single component per cell type (see Methods). Additionally, we considered the scenario of cell counts imputation wherein cell counts were known for 5% of the samples (*n* = 15; BayesCCE imp), and the scenario wherein samples from external data with both methylation levels and cell counts were used in the analysis (*n* = 15; BayesCCE imp ext). Top panel: mean absolute correlation (MAC) across all cell types. Bottom panel: mean absolute error (MAE) across all cell types. For BayesCCE imp and BayesCCE imp ext, the MAC and MAE values were calculated while excluding the samples with assumed known cell counts.

### BayesCCE impute: cell counts imputation

We next considered a scenario in which cell counts are known for a small subset of the samples in the data. This problem can be viewed as a problem of imputing missing cell count values (see Methods). We repeated all previous experiments, only this time we assumed that cell counts are known for randomly selected 5% of the samples in each data set. As opposed to the previous experiments, in which each one of the BayesCCE components constituted a scaled estimate of the proportions of one of the cell types, incorporating samples with known cell counts allowed BayesCCE to produce components that form absolute estimates of the cell type proportions (i.e. not scaled components, but components with low absolute error compared with the true proportions). Moreover, in contrast to previous experiments, each component was now automatically assigned to its corresponding cell type.

Under the assumption of six constituting cell types in blood tissue (*k* = 6), we observed a substantial improvement of up to 58% in mean absolute correlation values compared with our previous experiments (Figure 3 and Supplementary Tables ?? and ??). Specifically, we found the mean absolute correlation values across all six cell types to be 0.71, 0.66, 0.56 and 0.71 in the Hannum et al., Liu et al., Hannon et al. I and Hannon et al. II data sets, respectively. In addition, in contrast to our previous experiments, inclusion of some cell counts resulted in low mean absolute error, which reflects a correct scale for the components. We observed similar results when assuming three constituting cell types (*k* = 3), providing an improvement of up to 28% in correlation and a substantial decrease in absolute errors compared with the previous experiments (Supplementary Figure ?? and Supplementary Tables ?? and ??).

**Figure 3:**
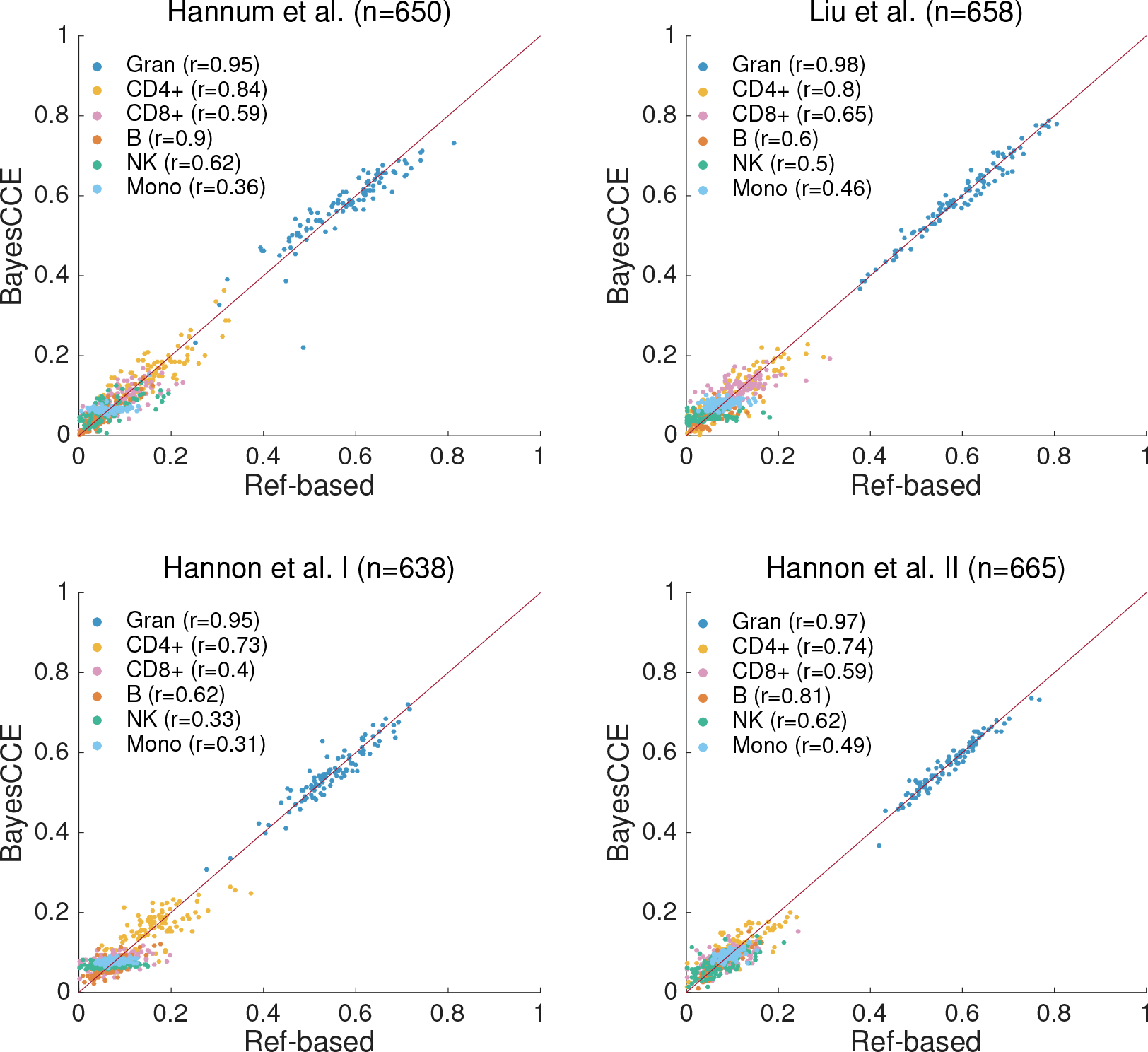
BayesCCE captures cell type proportions in four data sets under the assumption of six constituting cell types in blood (*k* = 6): granulocytes, monocytes and four subtypes of lymphocytes (CD4+, CD8+, B cells and NK cells), and assuming known cell counts for randomly selected 5% of the samples in the data. All correlations were calculated while excluding the samples with assumed known cell counts. For convenience of visualization, we only plot the results of 100 randomly selected samples for each data set.

In the absence of cell counts for a subset of the individuals in the data, we can incorporate into the analysis external data of samples for which both cell counts and methylation levels (from the same tissue) are available. We repeated again all previous experiments (*k* = 3 and *k* = 6), only this time for each data set we added a randomly selected subset of samples from one of the other data sets (5% of the original sample size), and used both their methylation levels and cell type proportions in the analysis. Specifically, we used randomly selected samples and corresponding estimates of cell type proportions from the Hannon et al. I data set for the experiments in all three other data sets, and samples from the Hannon et al. II data set for the experiment with the Hannon et al. I data set. In order to pool samples from two data sets together, we considered only the intersection of CpG sites that were available for analysis in the two data sets. In addition, unlike in the previous experiments, here we potentially introduce new batch effects into the analysis, as in each experiment the original sample is combined with external data. We therefore accounted for the new batch information by adding it as a new covariate into BayesCCE. As in the case of known cell counts for a subset of the samples, we found that the inclusion of external samples with both methylation and cell counts substantially improved the performance in terms of correlation and absolute errors (Supplementary Figures ?? and ?? and Supplementary Tables ?? and ??). These results clearly show that estimates can be dramatically more accurate given measured cell counts for as few as a couple of dozens of samples in the data (or such samples from external data).

As before, for assessing performance more thoroughly, we applied BayesCCE on the same sub-sampled data sets we used before (*n* = 300), while assuming known cell counts for a subset of the samples. In one scenario we assumed cell counts are known for 5% of the samples in each data set (*n* = 15), and in a second scenario we included into the analysis methylation levels and cell type proportions of 15 samples from external data. These experiments revealed in most cases a substantial improvement in correlation over a standard execution of BayesCCE (i.e. without inclusion of cell counts), and revealed in all cases a substantial improvement in mean absolute error. The results are summarized in Figure 2 for the case of six constituting cell types (*k* = 6) and in Supplementary Figure ?? for the case of three constituting cell types (*k* = 3).

We further tested the performance of BayesCCE as a function of the number of samples for which cell counts are available. Remarkably, we found that known cell counts for only a couple of dozens of the samples are needed in order to achieve the maximal improvement in performance; including more samples with known cell counts did not provide a further improvement (Figure 4). In addition, we evaluated the performance of BayesCCE as a function of the sample size. Interestingly, while performance did not improve by increasing the sample over a few hundred of samples in the case of unknown cell counts, we found that knowledge of cell counts for as few as 15 samples in the data allowed a monotonic improvement in performance in larger sample sizes (Supplementary Figure ??).

**Figure 4:**
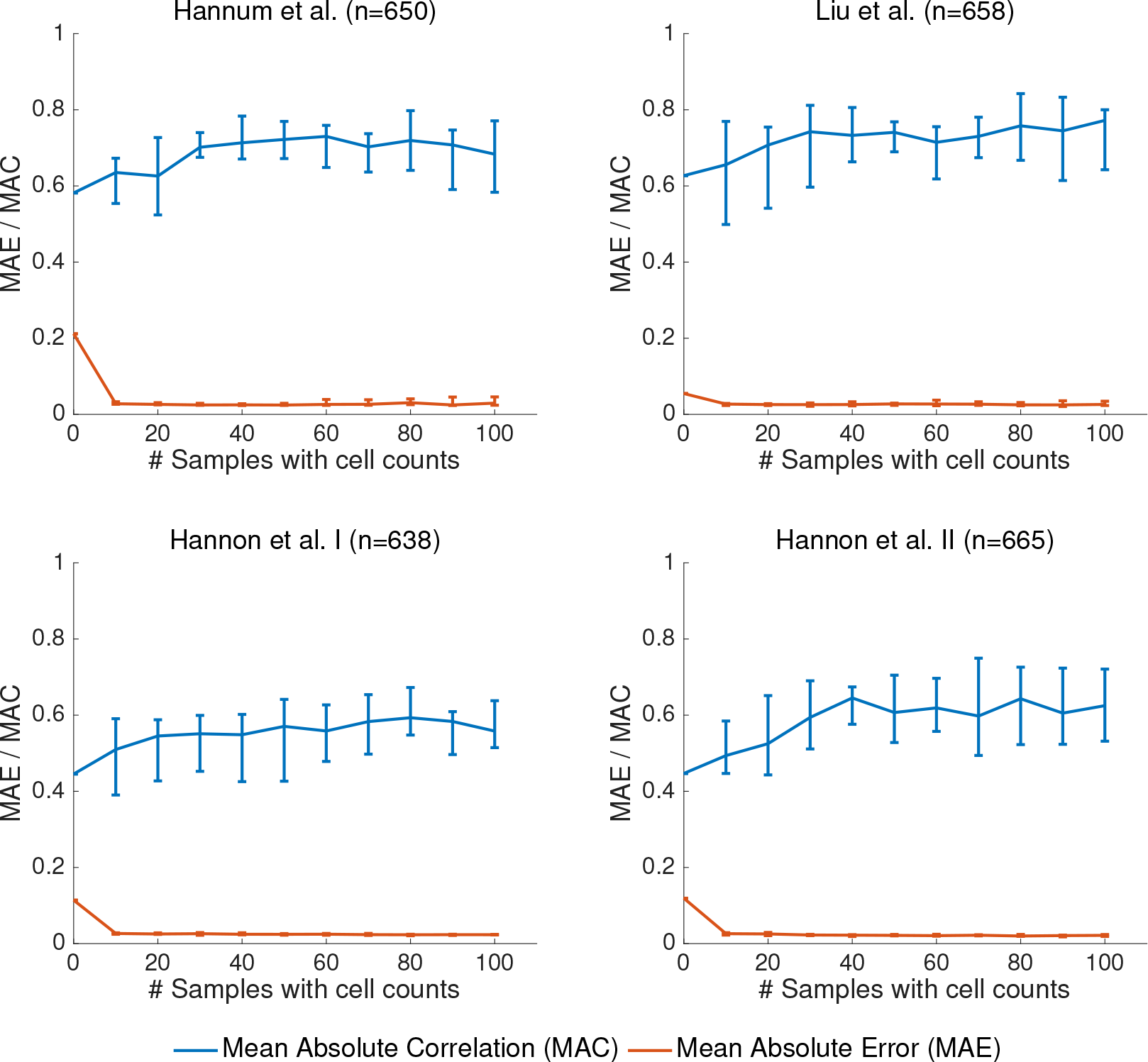
Performance of BayesCCE as a function of the number of samples for which cell counts are known, under the assumption of six constituting cell types in blood (*k* = 6): granulocytes, monocytes and four subtypes of lymphocytes (CD4+, CD8+, B cells and NK cells). Presented are the medians of the mean absolute correlation values (MAC; in blue) and the medians of the mean absolute error values (MAE; in red) across the six cell types. Error bars indicate the range of MAC and MAE values across ten different executions for each number of samples with known cell counts. In every execution samples with known cell counts were randomly selected, and all MAC and MAE values were calculated while excluding the samples with assumed known cell counts.

Finally, we considered an alternative approach for verifying the results of BayesCCE. Although our study aims at estimating cell-type proportions without the need for reference methylation data, BayesCCE jointly learns cell type composition and cell-type-specific mean methylation levels (methylomes). Hence, as a by-product of the BayesCCE algorithm, we also obtain cell-type-specific methylomes across the CpG sites selected by BayesCCE as part of its feature selection process (see Methods). Our experiments found BayesCCE to provide one component per cell type, however, these components are not necessarily appropriately scaled, which implies that estimated cell-type-specific methylation profiles are also not necessarily calibrated. Nevertheless, in the scenario where cell counts were known even for a small subset of the individuals in the study, BayesCCE provided calibrated cell count estimates. In such cases, we therefore expect BayesCCE to provide calibrated cell-type-specific methylation profiles. Using correlation maps, for each of the four whole-blood methylation data sets we analyzed, we verified high similarity between the cell-type-specific methylomes obtained by BayesCCE to those estimated by a reference methylation data collected from sorted blood cells [11] (Supplementary Figure ??). In spite of an overall high similarity between these two approaches, the correlation patterns detected by BayesCCE did not perfectly match those estimated using the reference data. While this may demonstrate the expected accuracy limitations of BayesCCE to some extent, we also attribute these imperfect matches, at least in part, to inaccuracies introduced by the reference data set, owing to the fact that it was constructed only from a small group of individuals (n = 6), which do not represent well all the individuals in other data sets in terms of methylome altering factors such as age [14], gender [15, 16], and genetics [17].

### Robustness of BayesCCE to biases introduced by the cell composition prior

BayesCCE relies on prior information about the distribution of the cell type composition in the studied tissue. In practice, the available prior information may not always precisely reflect the cell composition distribution of the individuals in the study. For instance, in a case/control study design, cases may demonstrate altered cell compositions compared with healthy individuals. Therefore, in this scenario, a prior estimated from a healthy population (or a sick population) is expected to deviate from the actual distribution in the sample. This potential problem is clearly not limited to case/control studies, but also applies to studies with quantitative phenotypes, in case these are correlated with changes in cell composition of the studied tissue. In principle, we can address this issue by incorporating several appropriate priors and assigning different priors to different individuals in the study. However, in practice, population-specific priors may be hard to obtain, mainly owing to the fact that numerous known and unknown factors can affect cell composition.

We revisited our analysis from the previous subsections in attempt to assess the robustness of BayesCCE to non-informative or misspecified priors. A desired behavior would allow BayesCCE to overcome a bias introduced by a prior which does not accurately represent all the individuals in the sample. Particularly, we considered three whole-blood case/control data sets, two schizopherenia data sets by Hannon et al. and a rheumatoid arthritis data set by Liu et al., all of which are expected to demonstrate differences in blood cell composition between cases and controls [26, 27]. In fact, in our analysis we had an inherently misspecified prior since we learned the prior from hospital patients (outpatients), which are overall expected to represent a sick population better than a more general population. Specifically, out of the 595 individuals used for learning the prior, 64% are known to have taken at least one medication at the time of blood draw for cell counting and 24% were admitted to the hospital due to various conditions within two months before or after the time of their blood draw (70.4% were either admitted or took medications). We expect these conditions to be correlated with alterations in blood cell composition, and therefore the prior information we used is expected to represent deviation from a healthy population and, as a result, to misrepresent at least the control individuals in the case/control data sets we analyzed. We further considered an additional fourth data set by Hannum et al., which was originally studied in the context of aging (age range: 19-101, mean: 64.03, SD: 14.73). Our prior was calculated using sample with a different distribution of ages (range: 20-88, mean: 49.19, SD: 16.69), thus potentially misrepresenting the cell composition distribution in the Hannum et al. data to some extent.

Remarkably, we found the cell composition estimates given by BayesCCE to effectively detect differences between populations in the data sets, in spite of using a single prior estimated from one particular population. Specifically, we found that BayesCCE correctly detected the cell types which differentiate between cases and controls and between young and older populations; notably, in some of the data sets we found BayesCCE to demonstrate some differences between cases and controls which were not captured by the reference-based estimates (Figure 5). For example, NK cells abundance is known to change in aging in a process known as NK cell immunesenescence [28, 29], and monocyte levels are known to increase in RA patients compared with healthy individuals. [30, 31, 32]. These differences in cell populations were detected by BayesCCE but not by the reference-based method, thus suggesting that BayesCCE could uncover signal which was detected by the reference-based method (Figure 5). We further estimated, for each data set, the distribution of white blood cells based on the BayesCCE cell count estimates, and verified the ability of BayesCCE to correctly capture two distinct distributions (cases and controls or young and older individuals), regardless of the single distribution encoded by the prior information (Figure 5). While BayesCCE provides one component per cell type, these components are not necessarily appropriately scaled to provide cell count estimates in absolute terms. Therefore, for the latter analysis, we considered only the scenarios in which cell counts are known for a small number of individuals.

**Figure 5:**
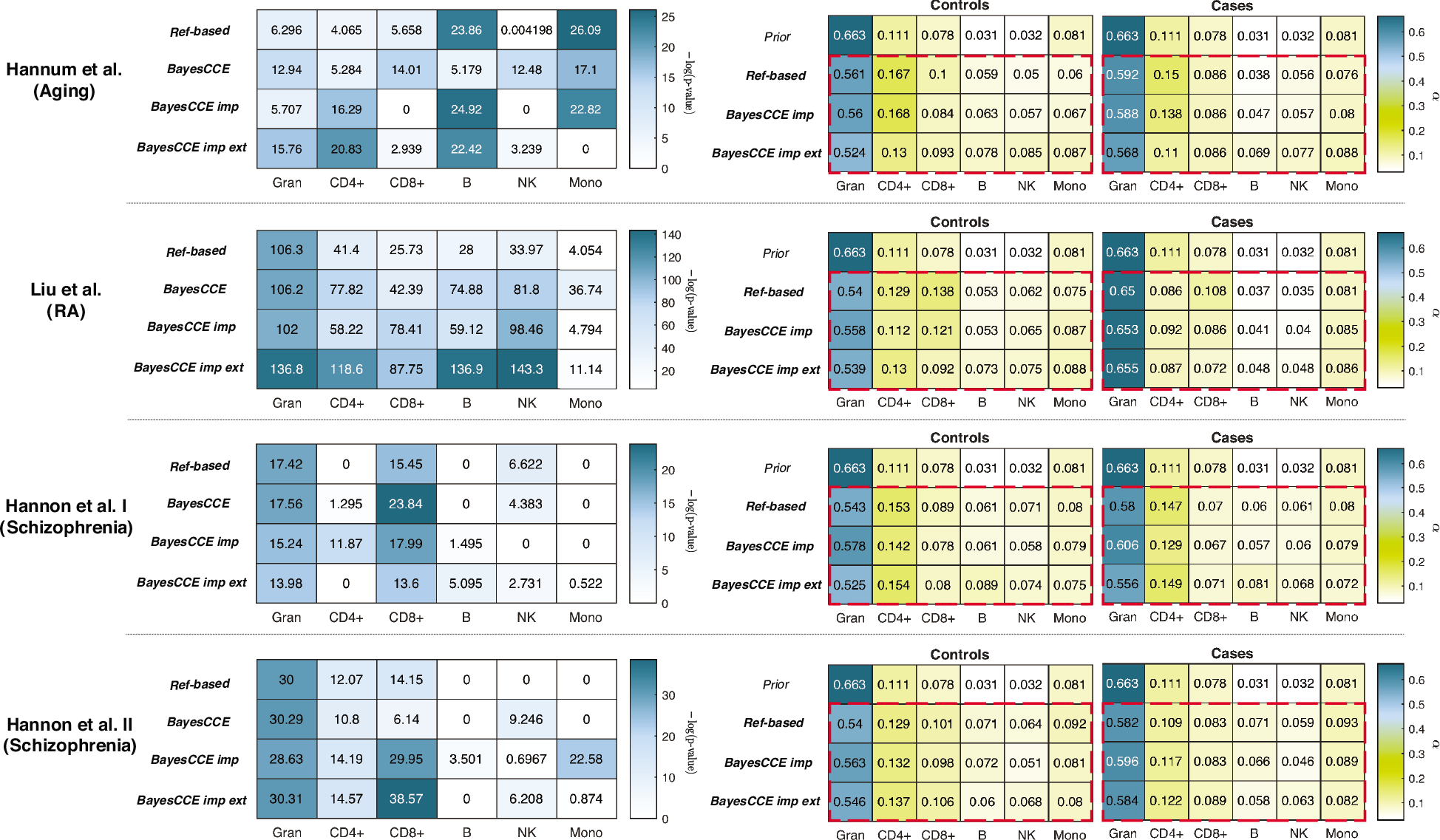
The robustness of BayesCCE to prior misspecification and its ability to capture population-specific variability in cell type composition. Considering six constituting cell types in blood (*k* = 6), granulocytes, monocytes and four subtypes of lymphocytes (CD4+, CD8+, B cells and NK cells), for each data set, presented are the negative log Bonferroni-adjusted p-values of a t-test between cases and controls using estimates of each cell type (left side), and the Dirichlet parameters of estimated cell counts stratified by cases and controls (right side). Results are presented for the cell count estimates obtained by the reference-based method, BayesCCE, BayesCCE with known cell counts for 5% of the samples (BayesCCE imp), and BayesCCE with 5% additional samples with both known cell counts and methylation from external data (BayesCCE imp ext). Red dashed rectangles emphasize the high similarity in estimated population-specific cell composition distributions between the different methods, regardless of the prior used (”prior”). For the Hannum et al. data set, cases were defined as individuals with age above the median age in the study. In the evaluation of BayesCCE imp and BayesCCE imp ext, samples with assumed known cell counts were excluded before calculating p-values and fitting the Dirichlet parameters.

We further evaluated the scenario in which two different population-specific prior distributions are available. Specifically, one prior for cases and another one for controls in the case/control studies, and one for young and another one for older individuals in the aging study. For the purpose of this experiment, we estimated the priors using the reference-based estimates of a subset of the individuals (5% of the sample size) that were then excluded from the rest of the analysis. Interestingly, we found the inclusion of two prior distributions to provide no clear improvement over using a single general prior (Supplementary Table ??). Thus, further confirming the robustness of BayesCCE to inaccuracies introduced by the prior information due to cell composition differences between populations.

Finally, we evaluated the effect of incorporating noisy priors on the performance of BayesCCE by considering a range of possible priors with different levels of inaccuracies, including a non-informative prior (Supplementary figure ??). Not surprisingly, we observed that given cell counts for a small subset of samples, BayesCCE was overall robust to prior misspecification, which did not result in a substantially reduced performance even given a non-informative prior. In the absence of known cell counts, the performance of BayesCCE was somewhat decreased, however, remained reasonable even in the scenario of a non-informative prior. Particularly, overall, BayesCCE with a non-informative prior performed better than the competing reference-free methods (ReFACTor, NNMF, and MeDeCom). We attribute this result to the combination of the constraints defined in BayesCCE with the sparse low-rank assumption it takes, which seems to handle more efficiently with the high-dimension nature of the computational problem (see Methods). We note that in the presence of a non-informative prior, BayesCCE conceptually reduces to the performance of ReFACTor, and therefore it captures the same cell composition variability in the data. Yet, owing to the additional constrains, BayesCCE allows to overcome ReFACTor in capturing a set of components such that each component corresponds to one cell type.

## 3 Discussion

We introduce BayesCCE, a Bayesian method for estimating cell type composition from heterogeneous methylation data without the need for methylation reference. We show mathematically and empirically the non-identifiability nature of the more straightforward reference-free NNMF approach for inferring cell counts, which tends to provide only linear combinations of the cell counts. In contrast, while we do not provide conditions for the uniqueness of a BayesCCE solution, our empirical evidence from multiple data sets clearly demonstrates the success of BayesCCE in providing desirable results of one component per cell type by leveraging readily obtainable priors from previously collected cell counts.

The parameters of the prior required for BayesCCE can be estimated by utilizing previous studies that collected cell counts from the tissue of interest. Since no other genomic information is required, obtaining such data is relatively easy for many tissues, such as brain [39], heart [40] and adipose tissue [41]. Particularly, such data should be substantially easier to obtain compared to reference data from sorted cells for the corresponding tissues. Ideally, in order to learn the prior, one would want to use cell counts coming from the same population as the target population. Nevertheless, empirically, we observe that BayesCCE leverages the prior to direct the solution while still allowing enough flexibility, which makes it robust even to substantial deviations of the prior from the true underlying cell composition distribution. In fact, our results demonstrate that BayesCCE handles biases introduced by the prior remarkably well. Particularly, it allows to capture differences in cell compositions between different populations in the same study, thus providing an opportunity to study cell composition differences between different populations even in the absence of methylation reference.

Since no large data sets with measured cell counts are currently publicly available, we used a supervised method [5] for obtaining cell type proportion estimates, which were used as the ground truth in our experiments. Even though the method used for obtaining these estimates was shown to reasonably estimate leukocyte cell proportions from whole blood methylation data in several independent studies [23, 18, 24], these estimates may have introduced biases into the analysis. Particularly, any inaccuracies introduced by the reference-based method could have directly affect the results of our evaluation. Our results indicate that such inaccuracies are more likely in some particular cell types over others. Failing to accurately estimate a particular cell type may be the outcome of various reasons. Notably, utilizing inappropriate reference data or failing to select a set of informative features that mark a particular cell type may dramatically affect its estimated values. Other reasons which are not methodological may also lead to inaccuracies of the estimates. For example, two cell types with very similar methylation patterns will be hardly distinguishable. In spite of the potential pitfalls of using estimates as a baseline for evaluation, we believe that our results on several independent data sets, including simulated data, and the use of a prior estimated from a large data set of high resolution cell counts, provide a compelling evidence for the utility of BayesCCE.

We further demonstrate that imputation of cell counts can be highly accurate when cell counts are available for some of the samples in the data. Particularly, based on our experiments, only as few as a couple of dozens of samples with known cell counts are needed in order to substantially improve performance. Moreover, in the general setup of BayesCCE, where no cell counts are known, each component corresponds to one cell type, however, not necessarily in the right scale and there is no automatic way to determine the identity of that cell type. In contrast, in the case of cell counts imputation, where cell counts are known for a subset of the samples, the assignment of components into cell types is straightforward. In addition, as we showed, BayesCCE is able to reconstruct cell counts up to a small absolute error (i.e. each component is scaled to form cell proportion estimates of one particular known cell type).

We note that in our evaluation of BayesCCE we considered only whole-blood data sets. Studying other tissues or biological conditions is clearly of interest. However, in the absence of other tissue-specific methylation references that were clearly shown to allow obtaining reasonable cell type proportion estimates, evaluation of performance based on tissues other than whole-blood will not be reliable. We therefore opt to focus on evaluating the performance of BayesCCE using multiple large whole-blood data sets. Importantly, beyond its potential utility for complex biological scenarios in which reference data is unavailable, BayesCCE may also provide an opportunity to improve cell count estimates in whole-blood studies in scenarios where the currently available reference data is not appropriate. Notably, in a recent work we have shown using multiple whole-blood data sets that ReFACTor outperforms the reference-based method in correcting for cell composition [19]. Differences in performance between ReFACTor (upon which BayesCCE relies for obtaining a starting point that captures the cell composition variation in the data) and the reference-based method are expected to be especially large in studies where the available reference data do not represent the individuals in the study well. We argue that this is likely to typically be the case, as the current go-to whole blood reference consists of only six individuals [11], which represent a very specific and narrow population in terms of methylome altering factors, such as age [14], gender [15, 16], and genetics [17]. That said, large data sets with experimentally measured cell counts are required in order to fully investigate and demonstrate these claims.

We further note that in our benchmarking of BayesCCE with existing reference-free methods we considered only a subset of the available methods in the literature. Other reference-free methods that have been suggested in the context of accounting for cell composition in methylation data exist, however, these do not provide explicit components, but rather only implicitly account for cell composition variability in association studies. While in principle these methods can be modified to produce components, in this work we focused only on methods that can be readily used to provide explicit components for evaluation. We further note that several supervised and unsupervised decomposition methods have been suggested for estimating cell composition from gene expression [34, 35, 36, 37, 38]. However, these were refined for gene expression data and, to the best of our knowledge, none of these methods takes into account prior knowledge about the cell composition distribution as in BayesCCE. It remains of interest to investigate whether BayesCCE can be adapted for estimating cell composition from gene expression without the need for purified expression profiles.

Finally, our approach is based on finding a suitable linear transformation of the components found by ReFACTor [8]. It is therefore important to follow the guidelines for the application of ReFACTor, such as incorporation of methylation altering covariates; these guidelines were recently highlighted elsewhere [33, 19]. Since BayesCCE relies on the ReFACTor components, it is limited by their quality, and particularly, if the variability of some cell type is not captured by ReFACTor, BayesCCE will not be able to estimate that cell type well. Such a result is possible in scenarios where the variation of a particular cell type is substantially weaker than other sources of variation in the data (which are unrelated to cell type composition); we note, however, that this potential limitation is not exclusive for ReFACTor or BayesCCE but rather a general limitation of all existing reference-free methods. BayesCCE will effectively provide the same result as ReFACTor if used for correcting for a potential cell type composition confounder in methylation data. Since ReFACTor does not allow to infer direct cell count estimates but rather linear transformations of those, we suggest to use BayesCCE in cases in which a study of individual cell types is performed and therefore ReFACTor cannot be used. In case merely a correction for cell composition is desired, we suggest to use BayesCCE when cell counts are known for a subset of the samples, and otherwise to use ReFACTor.

## 4 Conclusions

We introduce a Bayesian method for estimating cell type composition from heterogeneous methylation data using a prior on the cell composition distribution. In contrast to previous methods, using BayesCCE we can generate components such that each component corresponds to a single cell type. These components can allow researchers to perform types of downstream analyses that are not possible using previous reference-free methods, which essentially capture linear combinations of several cell types in each component they provide. Based on our results, showing a further substantial improvement by incorporating some cell counts into the analysis, we recommend that in future studies either the cell counts be measured for at least a couple of dozens of the samples or external data of samples with measured cell counts be utilized.

## 5 Methods

### Notations and related work

Let *O* ∈ ℝ^*m*×*n*^ be an m sites by *n* samples matrix of DNA methylation levels coming from a heterogeneous source consisting *k* cell types. For methylation levels, we consider what is commonly referred to as beta-normalized methylation levels, which are defined for each sample in each site as the proportion of methylated probes out of the total number of probes. Put differently, *O_ji_* ∈ [0,1] for each site *j* and sample *i*. We denote *M* ∈ ℝ^*m*×*k*^ as the cell-type-specific mean methylation levels for each site, and denote a row of this matrix, corresponding to the *j*th site, using *M*_*j*_,‥ Additionally, we denote *R* ∈ ℝ^*n*×*k*^ as the cell type proportions of the samples in the data. A common model for observed mixtures of DNA methylation is

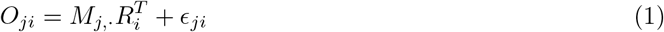

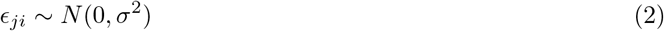

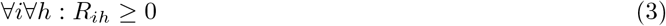

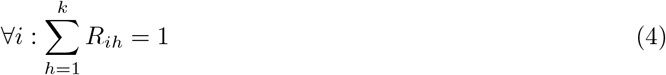

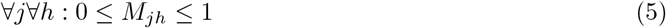

where the error term *∊*_*ji*_ models measurement noise and other possible unmodeled factors. The constraints in (3) and in (4) require the cell proportions to be positive and to sum up to one in each sample, and the constraints in (5) require the cell-type-specific mean levels to be in the range [0,1]. This model was initially suggested for DNA methylation in the context of reference-based estimation of cell proportions by Houseman et al. [5]. We are interested in estimating *R*. Taking a standard maximum-likelihood approach for fitting the model results in the following optimization problem:

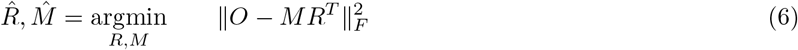

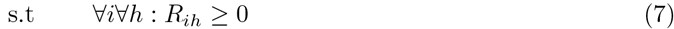

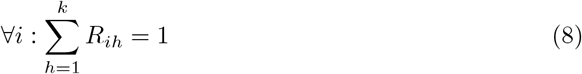

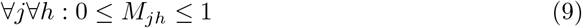

where 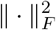 is the squared Frobenius norm. The reference-based method [5] first obtains an estimate of *M* from reference methylation data collected from sorted cells of the cell types composing the studied tissue. Once an estimate of *M* is fixed, *R* can be estimated by solving a standard quadratic program.

If the matrix *M* is unknown, which is a reference-free version of the problem, the above formulation of the problem can be regarded as a version of non-negative matrix factorization (NNMF) problem. NNMF has been suggested in several applications in biology; notably, the problem of inference of cell type composition from methylation data has been recently formulated as an NNMF problem [9]. In order to optimize the model, the authors used an alternative optimization procedure in which *M* or *R* are optimized while the other is kept fixed. However, as demonstrated by the authors [9], this solution results in the inference of a linear combination of the cell proportions *R*. Put differently, more than one component of the NNMF is required for explaining each cell type in the data. This was recently further highlighted and explained using geometric considerations [10], which nicely showed the non-identifiable nature of the NNMF model in (6) in case that a perfect factorization of *O* into *M*, *R* exists (i.e. *O* = *MR^T^*). However, in practice, perfect factorization never exists in real biological data. Thus, in addition to empirical evidence from several data sets on which we apply the NNMF method (see Results), in the next subsection we provide a mathematical proof for the non-identifiability of the NNMF model in (6) under a more general case, where a perfect factorization does not necessarily exist.

In an attempt to overcome the non-identifiability of the model in (6) and to provide cell type proportions when reference methylation data are not available, a recent modification of the NNMF model has been suggested [10]. The method, MeDeCom, solves the optimization of the NNMF model while including additional penalty term in the objective function. Derived from biological knowledge about mean methylation levels, the penalty negatively weights mean methylation levels diverging from a known bimodal behavior of methylation levels, wherein CpGs tend to be overall methylated or unmethylated [10]. While the modified objective suggested in MeDeCom overcomes the non-identifiability of the NNMF model for a given weight of the penalty (*λ*), it is not entirely clear how to select *λ*. To circumvent this problem, the authors proposed a cross-validation procedure for the selection of *λ*. However, our empirical results from four large whole-blood methylation data sets, as well as from simulated data, show sub-optimal performance for MeDeCom, similarly to the solutions of the simpler NNMF model. Our results suggest that the modification introduced by MeDeCom may not effectively avoid the non-identifiability nature of the NNMF model, possibly due to insufficient prior information or inability to effectively determine an appropriate value for *λ*.

Another recent reference-free method for estimating cell composition in methylation data, ReFACTor [8], performs an unsupervised feature selection step followed by a principal components analysis (PCA). Similarly to the NNMF solution, ReFACTor is an unsupervised method and it only finds principal components (PCs) that form linear combinations of the cell proportions rather than directly estimates the cell proportion values [8].

### Non-identifiability of the NNMF model

We hereby show by construction the non-identifiability nature of the NNMF model in (6). For this proof, instead of the constraints in (9), we consider a slightly modified version of the constraints:

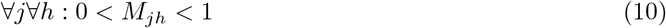

While in theory we may have an equality (i.e. *M_jh_* = 0 or *M_jh_* = 1), in practice, such sites are typically not measured or excluded from the analysis, since they would not be demonstrating any variability.

#### Proposition

Let 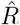, 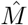 be a solution to the problem in (6). There exist 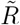 ≠ 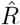, 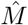 ≠ 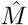 such that 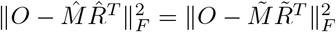 and the constraints in (7), (8) and in (10) are satisfied.

#### Proof

Let 0 > *c* > 1, define *Q* ∈ ℝ^*k*×*k*^ to be the identity matrix up to two entries: *Q*_11_ = 1 — *c,Q*_12_ = *c*. It follows that *Q*^−1^ is also the identity matrix up to two entries: 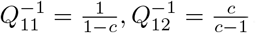.

Denote 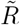 = 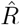*Q* and denote 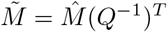, we get that

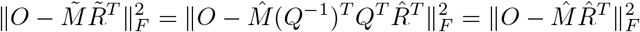

The constraints in (7) hold since 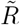_*ih*_ ≥ 0 for each 1 ≤ *i* ≤ *n*, 1 ≤ *h* ≤ *k*. The constraints in (8) hold since for each 1 ≤ *i* ≤ *n*

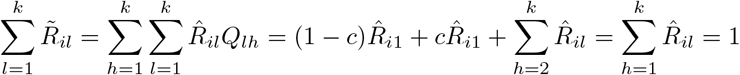

In addition, 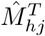 ∈ (0,1) for 2 ≤ *h* ≤ *k*, 1 ≤ *j* ≤ *m*. In order to completely satisfy the constraints in (10), we also require these constraints to be satisfied for *h* =1, 1 ≤ *j* ≤ *m*. It is easy to see that for each *j* the latter is satisfied if

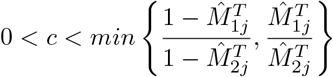

Therefore, we can simply select a value of *c* in the range

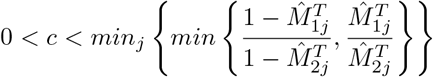

Note that we necessarily either have

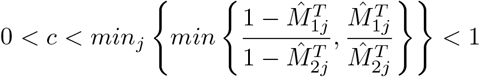

or

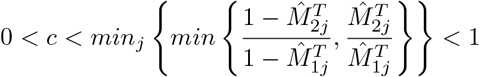

In the latter case we can switch the positions of the first two columns in *M*. Equality of the minimum to 1 in both cases would mean that *M*_1_ = *M*_2_, which would mean that the problem is non-identifiable, as the first two cell types cannot be distinguished in this scenario. As a result of the above, the constraints in (10) can be satisfied for a range of values of *c*. ⁊

### Model

We suggest a more detailed model by adding a prior on *R* and taking into account potential covariates. Specifically, we assume that

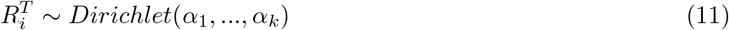

where *α*,…, *α_k_* are assumed to be known. In practice, the parameters are estimated from external data in which cell type proportions of the studied tissue are known. Such experimentally obtained cell type proportions were used to test the appropriateness of the Dirichlet prior in describing cell composition distribution (data not shown). Also, we consider additional factors of variation affecting observed methylation levels, in addition to variation in cell type composition. Specifically, denote *X* ∈ ℝ^*n*×*p*^ as a matrix of *p* covariates for each individual and *S* ∈ ℝ^*m*×*p*^ as a matrix of corresponding effects of the *p* covariates on each of the *m* sites. As before, we are interested in estimating *R*. Deriving a maximum likelihood-based solution for this model and repeating the constraints for completeness results in the following optimization problem:

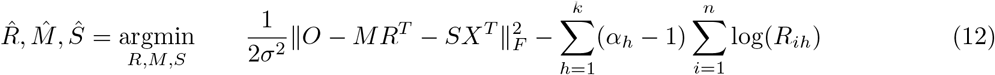

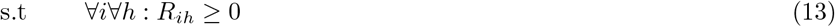

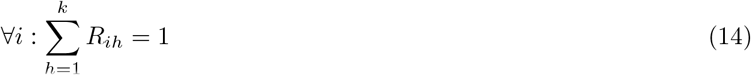

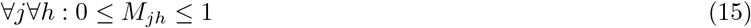

Our intuition in this model is that since the priors on *R* are estimated from real data, incorporating them will push the solution of the optimization to return estimates of *R* which are closer to the true values as opposed to a linear combination of them.

### Algorithm

Our algorithm uses ReFACTor as a starting point. Specifically, we estimate *R* by finding an appropriate linear transformation of the ReFACTor principal components (ReFACTor components). In principle, any of the reference-free methods we examined (ReFACTor, NNMF and MeDeCom) could be used as the starting point for our method. However we found that ReFACTor captures a larger portion of the cell composition variance compared with the alternatives (Supplementary Figure ??).

Applying ReFACTor on our input matrix *O* we get a list of *t* sites that are expected to be most informative with respect to the cell composition in *O*. Let 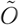 ∈ ℝ^*t*×*n*^ be a truncated version of *O* containing only the *t* sites selected by ReFACTor. We apply PCA on 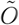 to get *L* ∈ ℝ^*t*×*d*^, *P* ∈ ℝ^*n*×*d*^, the loadings and scores of the first *d* ReFACTor components. Then, we reformulate the original optimization problem in terms of linear transformations of *L* and *P* as follows:

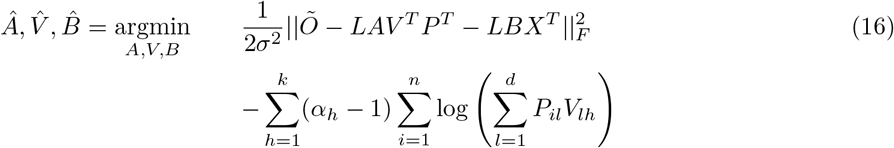

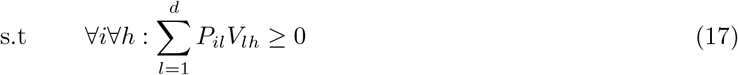

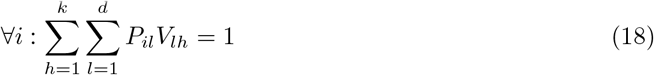

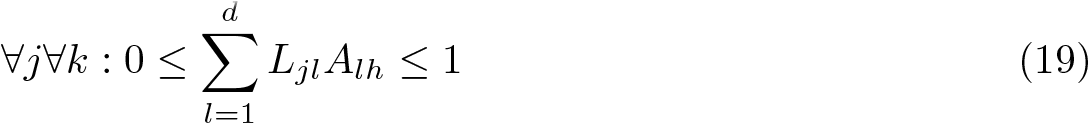

where *A* ∈ ℝ^*d*×*k*^ is a transformation matrix such that 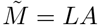 (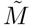 being a truncated version of *M* with the *t* sites selected by ReFACTor), *V* ∈ ℝ^*d*×*k*^ is a transformation matrix such that *R* = *PV* and *B* ∈ ℝ^*d*×*p*^ is a transformation matrix such that *LB* corresponds to the effects of each covariate on the methylation levels in each site. The constraints in (17) and in (18) correspond to the constraints in (13) and in (14), and the constraints in (19) correspond to the constraints in (15).

Given 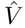, we simply return 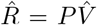 as the estimated cell proportions. Note that in the new formulation we are now required to learn only *d*(2*k* + *p*) parameters – *d, k* and *p* being small constants - a dramatically decreased number of parameters compared with the original problem which requires *nk* +*m*(*k*+*p*) parameters. By taking this approach, we make an assumption that 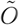 consists of a low rank structure that captures the cell composition using *d* orthogonal vectors. While a natural value for *d* would be *k, d* is not bounded to be *k*. Particularly, in cases where substantial additional cell composition signal is expected to be captured by latter ReFACTor components (i.e. components beyond the first *k*), we would expect to benefit from increasing *d*. Clearly, overly increasing d is expected to result in overfitting and thus a decrease in performance. Finally, taking into account covariates with potentially dominant effects in the data should alleviate the risk of introducing noise into 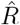 in case of mixed low rank structure of cell composition signal and other unwanted variation in the data. We note, however, that similarly to the case of correlated explaining variables in regression, considering covariates that are expected to be correlated with the cell type composition may result in underestimation of *A*, *V* and therefore to a decrease in the quality of 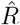.

### Imputing cell counts using a subset of samples with measured cell counts

In practice, we observe that each of BayesCCE’s components corresponds to a linear transformation of one cell type rather than to an estimate of that cell type in absolute terms. That is, it still lacks the right scaling (multiplication by a constant and addition of a constant) for transforming it into cell type proportions. Furthermore, we would like the *i*th BayesCCE component to correspond to the *i*th cell type described by the prior using the *α*_*i*_ parameter. Empirically, this is not necessarily the case, especially in scenarios where some of the *α*_*i*_ values are similar. In order to address these two caveats, we suggest incorporating measured cell counts for a subset of the samples in the data.

Assume we have *n*_0_ reference samples in the data with known cell counts*R*^(0)^ and *n*_1_ samples with unknown cell counts *R*^(1)^ (*n* = *n*_0_ + *n*_1_). This problem can be regarded as an imputation problem, in which we aim at imputing cell counts for samples with unknown cell counts. We can find 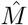 by solving the problem in (12) under the constraints in (15) for the *n*_0_ reference samples while replacing *R* with *R*^(0)^ and keeping it fixed. Then, given 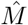, we can now solve the problem in (16), after replacing *LA* with 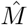 (i.e. we find only *V,B* now), under the following constraints

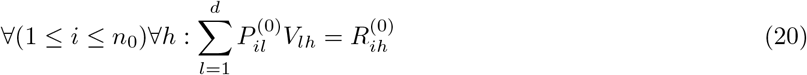

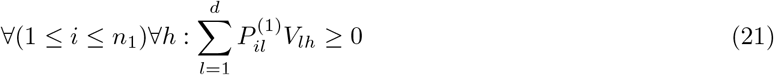

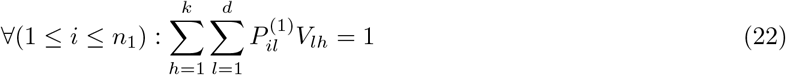

where *P*^(0)^ contains *n*_0_ rows corresponding to the reference samples in *P*, and *P*^(1)^ contains *n*_1_ rows corresponding to the remaining samples in *P*. In this case, both problems of estimating *M* and solving (16) while keeping *M* fixed are convex - the first problem takes the form of a standard quadratic problem and the latter results in an optimization problem of the sum of two convex terms under linear constraints. Using *M*, estimated from cell counts and corresponding methylation levels of a group of samples, as well as adding the constraints in (20), are expected to direct the inference of R towards a set of components such that each one corresponds to one known cell type with a proper scale.

### Implementation and practical issues

We estimate σ^2^ in (16) as the mean squared error of predicting 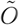 with *P* and *X*. The *α*_1_,…, *α*_*k*_ Dirichlet parameters of the prior can be estimated from cell counts using maximum likelihood estimators. In practice, we add a column of ones to both *L* and *P* in (16) in order to assure feasibility of the problem - these constant columns are used to compose the mean methylation level per site across all cell types and the mean cell proportion fraction in each cell type across all samples. In addition, we slightly relax some of the constraints in the problem to avoid problems due to numeric instability and inconsistent noise issues. First, the inequality constraints in (17) and in (21) are changed to require the cell proportions to be greater than *∊* > 0, as a result of the logarithm term in the objective (*∊* = 0.0001). In addition, we do not impose the equality constraints in (18) and in (22) but rather allow a small deviation from equality (5%), and given cell counts for a subset of the samples, we allow a small deviation from the equality constraints in (20) due to expected inaccuracies of cell count measurements (1%). The last two constraints are required for assuring a feasible solution, owing to the fact that we fit 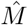, 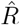 jointly. Specifically, since the starting point for the optimization is essentially a set of principal components (given by ReFACTor), which are not guaranteed to capture only cell composition variation, in practice, obtaining linear transformations that precisely satisfy the constraints is expected to be an exception rather than the rule. We verified this empirically (data not shown), and we further observed that these relaxations eventually result in feasible solutions which typically tend to tightly concentrate around the original constraints.

We performed all the experiments in this paper using a Matlab implementation of BayesCCE. Specifically, we solved the optimization problems in BayesCCE using the *fmincon* function with the default interior-point algorithm, and we used the *fastfit* [42] Matlab package for calculating maximum likelihood estimates of the Dirichlet priors. All executions of BayesCCE required less than an hour (and typically several minutes) on a 64-bit Mac OS X computer with 3.1GHz and 16GB of RAM. Corresponding code is available at: https://github.com/cozygene/bayescce.

### Evaluation of performance

The fraction of cell composition variation (*R*^2^) captured by each of the reference-free methods, ReFACTor, NNMF and MeDeCom, was computed for each cell type using a linear predictor fitted with the first *k* components provided by each method. In order to evaluate the performance of BayesCCE, for each component *i* we calculated its absolute correlation with the *i*th cell type, and reported the mean absolute correlation (MAC) across the *k* estimated cell types. While the Dirichlet prior assigns a specific parameter *α_h_* for each cell type *h*, empirically, we observed that in the case of *k* = 6 with no known cell counts for a subset of the samples, the *i*th BayesCCE component did not necessarily correspond to the *i*th cell type. Put differently, the labels of the *k* cell types had to be permuted before calculating the MAC. In this case we considered the permutation of the labels which resulted with the highest MAC as the correct permutation. In the rest of the cases, we did not apply such permutation (all the experiments using *k* = 3 and all the experiments using *k* = 6 with known cell counts for a subset of the samples).

For evaluating ReFACTor, NNMF and MeDeCom, reference-free methods which do not attribute their components to specific cell types in any scenario, we considered for each method the permutation of its components leading to the highest MAC in all experiments when compared with BayesCCE. In addition, we considered absolute error of the estimates from the ground truth as an additional quality measurement. We calculated the mean absolute error (MAE) across the k estimated cell types. When calculating absolute errors for the ReFACTor components, we scaled each ReFACTor component to be in the range [0, 1].

### Implementation and application of the reference-free and reference-based methods

We calculated the ReFACTor components for each data set using the parameters *k* = 6 and *t* = 500 and according to the default implementation and recommended guidelines of ReFACTor as described in the GLINT tool [33] and in a recent work [19], while accounting for known covariates in each data set. More specifically, in the Hannum et al. data [20] we accounted for age, sex, ethnicity and batch information, in the Liu et al. data [21] we accounted for age, sex, smoking status and batch information, and in the two Hannon et al. data sets [22] we accounted for age, sex and case / control state. We used the first six ReFACTor components (*d* = 6) for simulated data in order to accommodate with the number of simulated cell types, and the first ten components (*d* = 10) for real data, as real data are typically more complex and are therefore more likely to contain substantial signal in latter components.

The NNMF components were computed for each data set using the default setup of the RefFreeEWAS R package from the subset of 10,000 most variable sites in the data set, as performed in the NNMF paper by the authors [9]. Similarly, the MeDeCom components were computed for each data set using the default setup of the MeDeCom R package [10] from the subset of 10,000 most variable sites in the data set, as repeatedly running the method on the entire set of CpGs was revealed to be computationally prohibitive. The regularization parameter *λ* was selected according to a minimum cross-validation error criterion, as instructed in the MeDeCom package.

We used the GLINT tool [33] for estimating blood cell type proportions for each one of the data sets, according to the Houseman et al. method [5], using 300 highly informative methylation sites defined in a recent study [24] and using reference data collected from sorted blood cells [11].

### Data sets

We evaluated the performance of BayesCCE using a total of six data sets, as described bellow. For the real data experiments we downloaded four publicly available Illumina 450K DNA methylation array data sets from the Gene Expression Omnibus (GEO) database: a data set by Hannum et al. (accession GSE40279) from a study of aging rate [20], a data set by Liu et al. (accession GSE42861) from a recent association study of DNA methylation with rheumatoid arthritis [21], and two data sets by Hannon et al. (accessions GSE80417 and GSE84727; denote Hannon et al. I and Hannon et al. II) from a recent association study of DNA methylation with schizophrenia).

We preprocessed the data according to a recently suggested normalization pipeline [43]. Specifically, we retrieved and processed raw IDAT methylation files using R and the minfi R package [44] as follows. We removed 65 single nucleotide polymorphism (SNP) markers and applied the Illumina background correction to all intensity values, while separately analyzing probes coming from autosomal and non-autosomal chromosomes. We used a detection P-value threshold of P-value < 10^−16^ for intensity values, setting probes with P-values higher than this threshold to be missing values. Based on these missing values, we excluded samples with call rates < 95%. Since IDAT files were not made available for the Hannum et al. data set, we used the methylation intensity levels published by the authors.

As for data normalization, following the same suggested pipeline [43], we performed a quantile normalisation of the methylation intensity values, subdivided by probe type, probe sub-type and color channel. Beta normalized methylation levels were eventually calculated based on intensities levels (according to the recommendation by Illumina). On top of that, we excluded probes with over 10% missing values and used the “impute” R package for imputing remaining missing values. Additionally, using GLINT [33], we excluded from each data set all CpGs coming from the non-autosomal chromosomes, as well as polymorphic and cross-reactive sites, as was previously suggested [45].

We further removed outlier samples and samples with missing covariates. In more details, we removed six samples from the Hannum et al. data set and two samples from the Liu et al. data set, which demonstrated extreme values in their first two principal components (over four empirical standard deviations). Furthermore, we removed from the Liu et al. data set two additional remaining samples that were regarded as outliers in the original study of Liu et al., and we removed from the Hannon et al. data sets samples with missing age information. The final number of samples remained for analysis were *n* = 650, *n* = 658, *n* = 638 and *n* = 656, and the numbers of CpGs remained were 382,158, 376,021, 381,338 and 382,158, for the Hannum et al. data set, Liu et al. data set, and the Hannon et al. I and Hannon et al. II data sets, respectively.

For learning prior information about the distribution of blood cell type proportions we used electronic medical record (EMR) based study data that were acquired via the previously published Department of Anesthesiology and Perioperative Medicine at UCLA’s perioperative data warehouse (PDW) [46]. The PDW is a structured reporting schema that contains all the relevant clinical data entered into an EMR via the use of Clarity, the relational database created by EPIC (EPIC Systems, Verona, WI) for data analytics and reporting. We used high resolution cell counts measurements from adult individuals (*n* = 595) for fitting a Dirichlet distribution. The resulted parameters of the prior were 15.0727, 1.8439, 2.5392, 1.7934, 0.7240 and 0.7404 for granulocytes, monocytes, CD4+, CD8+, B cells, and NK cells, respectively. The parameters of the prior calculated for the case of three assumed cell types (*k* = 3) were 7.7681, 0.9503, and 2.9876 for granulocytes, monocytes, and lymphocytes, respectively. Finally, for generating simulated data sets and for generating correlation maps of cell-type-specific methylomes, we used publicly available data of methylation reference of sorted cell types collected in six individuals from whole blood tissue (GEO accession GSE35069) [11].

### Data simulation

We simulated data following a model that was previously described in details elsewhere [8]. Briefly, we used methylation levels from sorted blood cells [11] and, assuming normality, estimated maximum likelihood parameters for each site in each cell type. cell-type-specific DNA methylation data were then generated for each simulated individual from normal distributions with the estimated parameters, conditional on the range [0,1], for six cell types and for each site. Cell proportions for each individual were generated using a Dirichlet distribution with the same parameters used in the real data analysis. Eventually, observed DNA methylation levels were composed from the cell-type-specific methylation levels and cell proportions for each individual, and a random normal noise was added to every data entry to simulate technical noise (σ = 0.01). To simulate inaccuracies of the prior, the Dirichlet parameters required by BayesCCE were learned from cell type proportions of 50 samples generated at random from a Dirichlet distribution using the parameters learned from real data.

## Acknowledgments

This research was partially supported by the Edmond J. Safra Center for Bioinformatics at Tel Aviv University. E.H., E.R., L.S. and R.S. were supported in part by the Israel Science Foundation (Grant 1425/13), E.H., L.S. and R.S. by the United States Israel Binational Science Foundation grant 2012304. E.R. and L.S. were supported by Len Blavatnik and the Blavatnik Research Foundation. R.S. was supported by the Colton Family Foundation. E.E. was supported by National Science Foundation grants 0513612, 0731455, 0729049, 0916676, 1065276, 1302448, 1320589 and 1331176, and National Institutes of Health grants K25-HL080079, U01-DA024417, P01-HL30568, P01-HL28481, R01-GM083198, R01-ES021801, R01-MH101782 and R01-ES022282

